# miR-210 expression is strongly hypoxia-induced in anaplastic thyroid cancer cell lines and is associated with extracellular vesicles & Argonaute-2

**DOI:** 10.1101/2022.11.23.515840

**Authors:** Bonita H. Powell, Andrey Turchinovich, Yongchun Wang, Olesia Gololobova, Dominik Buschmann, Martha A. Zeiger, Christopher B. Umbricht, Kenneth W. Witwer

**Author notes:** **Address correspondence to:** Kenneth W. Witwer, PhD, 733. N. Broadway, Miller Research Building Room 827, Baltimore, MD 21205.

## Abstract

Hypoxia, or low oxygen tension, is frequently found in highly proliferative solid tumors such as anaplastic thyroid carcinoma (ATC) and is believed to promote resistance to chemotherapy and radiation. Identifying hypoxic cells for targeted therapy may thus be an effective approach to treating aggressive cancers. Here, we explore the potential of the well-known hypoxia-responsive microRNA (miRNA) miR-210-3p as a cellular and extracellular biological marker of hypoxia. We compare miRNA expression across several ATC and papillary thyroid cancer (PTC) cell lines. In the ATC cell line SW1736, miR-210-3p expression levels indicate hypoxia during exposure to low oxygen conditions (2% O_2_). Furthermore, when released by SW1736 cells into the extracellular space, miR-210-3p is associated with RNA carriers such as extracellular vesicles (EVs) and Argonaute-2 (AGO2), making it a potential extracellular marker for hypoxia.

## INTRODUCTION

Anaplastic thyroid carcinoma (ATC) is a rare malignancy, yet it accounts for the majority of all thyroid tumor-related deaths ^1–4^. In contrast with thyroid cancers such as papillary thyroid cancer (PTC), ATC cases are characteristically aggressive and almost always fatal, as these tumors acquire numerous deleterious genetic aberrations during de-differentiation ^1,5^. ATCs, like many other types of solid tumors, are highly proliferative and often hypoxic (oxygen-deprived)^5–8^. Tumors adapt to hypoxic stress by activating various cell signaling pathways through the action of hypoxia-inducible factors (HIFs). HIF-1 is a heterodimer composed of an alpha and a beta subunit^9^^10^. In normoxic conditions, HIF-1α is produced constitutively but is rapidly degraded by the proteasome upon hydroxylation. This occurs at specific proline residues by prolyl hydroxylase (PHD), followed by ubiquitination by the von Hippel-Lindau (VHL) protein^11–13^. In hypoxic conditions, HIF-1α proteins remain stable and translocate to the nucleus to activate genes by associating with the HIF-1ß subunit and binding to HIF Response Elements (HREs) within the proximal promoters of target genes^14–16^. Several HIF targets are involved in cell cycle regulation, metabolism, and angiogenesis^14,17–22^. Furthermore, hypoxic tumor cells with elevated HIF-1α protein levels are often resistant to chemotherapy and ionizing radiation^8,23,24^. Hence, developing approaches to identify and target hypoxic cells within ATC tumors may improve patient outcomes.

miRNAs are short, ∼22 nucleotide, post-transcriptional gene regulators that bind to complementary regions within the 3’UTR of mRNAs, thereby inhibiting translation or mediating degradation^25–28^. Cells release miRNAs into the extracellular milieu, where they are protected from RNase digestion in carriers including extracellular vesicles (EVs) and extracellular proteins such as Argonaute-2 (AGO2)^29–35^. Thus, miRNAs may be particularly attractive for the purpose of “liquid biopsies”: using biological fluids to assess the presence and state of cancer cells in the body. We and several others have reported dysregulated microRNAs (miRNAs) in thyroid cancers. The most commonly aberrantly expressed miRNAs in ATC include miR-222, miR-221, miR-146b and miR-21^36–41^. However, there is a paucity of information on miRNA expression differences in hypoxic vs. non-hypoxic regions within thyroid tumors ^30–35^.

In this study, we examined miR-210-3p (“miR-210”) as a potential marker of hypoxia. miR-210 is a well-known hypoxia-responsive miRNA and a direct HIF-1 target^42–45^. Its expression is frequently up-regulated in hypoxic tumor cells. Like HIF-1α protein levels, elevated miR-210 expression is associated with a poor prognosis^46,47^. We report herein on miR-210 expression in both PTC and ATC cell lines, examine the influence of hypoxia on cellular and extracellular miR-210-3p in the ATC line SW1736, and assess the extracellular association of miR-210 with EVs and AGO2.

## MATERIALS & METHODS

### Cell Culture

SW1736, C643, BCPAP and TPC-1 thyroid cell lines ^48–51^ were authenticated by the Johns Hopkins Genetic Resources Core Facility by Short Tandem Repeat (STR) profiling. Cells were cultured in RPMI-1640 medium (Gibco cat#11875-119) supplemented with: 10% Exosome-Depleted fetal bovine serum (Gibco A27208-01, Lot#2165597), 2mM L-glutamine (Thermo Fisher 25030-081), 1% Non-Essential Amino Acids (Thermo Fisher cat#1140050), 10mM HEPES (Life Tech cat#5630-080), 100 U/mL Penicillin Streptomycin (Thermo Fisher cat#15140-122). Cells were cultured in a humidified 37°C, 5% CO_2_ incubator under 21% O_2_ (normoxic) or in a 2% O_2_ (hypoxic) chamber, balanced with N_2_ using a compact O_2_/CO_2_ Controller ProOx c21 (Biospherix RS485) for 2, 4, 8, 24, 48, 72, or 96 hours.

### Cell Viability & Metabolic Activity Assays

Cell counts and viability were obtained using the Muse cell analyzer (EMD Millipore cat#1000-5175 serial#1846277) and Muse Viability reagent (Millipore cat#MCH600103). Glucose concentrations from conditioned media were measured using a glucose monitoring system (bioreactor sciences GM-100 serial#15100005) and test strips (GMTS-50 bioreactor science).

### Wound Healing Assay

Cells were seeded into 6-well plates and cultured until 100% confluent in normoxic conditions. Media was removed from wells and cells were washed twice gently with PBS. A 200µL pipette was used to create a scratch. Cells were washed again, and a sharpie was used to mark a location for imaging. Images were taken at 0hrs and 24hrs during hypoxia.

### EV Production and Separation

Cell culture conditioned medium was collected from cells and centrifuged at 1,000xg for 5min to pellet dead cells. The supernatant was centrifuged at 2,800xg for 10min to clear cell debris and apoptotic bodies. The 2,800xg supernatant was then concentrated down to 500µL at 3,500xg using Centricon Plus-70 10kD cut-off concentrators (Millipore cat#UFC701008) for 1hr at 4°C. 500µL of concentrated medium was fractionated over a qEV original 70nm size-exclusion chromatography column according to the manufacturer’s instructions (IZON Science SP1). Pooled EV fractions (7-9) were concentrated down to ∼100µL at 4,000xg using Amicon Ultra-2 Centrifugal Filter Units10kD cut-off concentrators (Millipore cat#UFC201024) at 4°C.

### Electron Microscopy

EV and protein fractions were negatively stained and analyzed by transmission electron microscopy (TEM) at the Johns Hopkins Microscope Core Facility as previously described^52^. Samples were adsorbed (10µL) to glow-discharged (EMS GloQube) carbon coated 400 mesh copper grids (EMS), by flotation for 2 min. Grids were quickly blotted then rinsed in 3 drops (1 min each) of tris-buffered-saline (TBS). Grids were negatively stained in 2 consecutive drops of 1% uranyl acetate with tylose (1% UAT, double filtered, 0.22 µm filter), blotted then quickly aspirated to get a thin layer of stain covering the sample. Grids were imaged on a Phillips CM-120 TEM operating at 80 kV with an AMT XR80 CCD (8 megapixel).

### Nano-flow Cytometry Measurement (NFCM)

The NanoFCM Flow NanoAnalyzer was used to measure concentration and particle size following the manufacturer’s instructions and as described previously ^52,53^. The instrument was calibrated separately for concentration and size using 250-nm Quality Control Nanospheres (NanoFCM) and a Silica Nanosphere Cocktail (NanoFCM cat#S16M-Exo) for detection of side scatter (SSC) of individual particles. Events were recorded for 1 min. Using the calibration curve, the flow rate and side scattering intensity were converted into corresponding particle concentrations and size.

### Single Particle Interferometric Reflectance Imaging (SP-IRIS)

Measurements were performed as described previously^52^. 50μl of EVs were diluted 1:1 in 1X Incubation Solution II and incubated at room temperature on ExoView Human Tetraspanin chips (Unchained Labs, Pleasanton, CA, Cat # 251-1000) printed with anti-human CD81 (JS-81), anti-human CD63 (H5C6), anti-human CD9 (HI9a), and anti-mouse IgG1 (MOPC-21). After incubation for 16hrs, chips were washed with 1X Solution A I 4 times for 3min each under gentle horizontal agitation at 460 rpm. Chips were then incubated for 1hr at room temperature with a fluorescent antibody cocktail of anti-human CD81 (JS-81, CF555), anti-human CD63 (H5C6, CF647), and anti-human CD9 (HI9a, CF488A) at a dilution of 1:1200 (v:v) in a 1:1 (v:v) mixture of 1X Solution A I and Blocking Solution II. The buffer was then exchanged to 1X Solution A I only, followed by 1 wash with 1X Solution A I, 3 washes with 1X Solution B I, and 1 wash with water (3min each at 460 rpm). Chips were immersed in water for approximately 10s each and removed at a 45-degree angle to allow the liquid to vacate the chip. All reagents and antibodies were supplied by Unchained Labs (Pleasanton, CA). Samples were diluted in 1X Incubation Solution II to load 50μL of 4.0×10^8 particles/mL, nominally, per chip. All chips were imaged in the ExoView R100 (Unchained Labs, Pleasanton, CA) by interferometric reflectance imaging and fluorescent detection. Data were analyzed using ExoView Analyzer 3.1 Software (Unchained Labs). Fluorescent cutoffs were as follows: CD63 channel 200, CD81 channel 400, CD9 channel 400.

### Cell and EV Lysis

Cells were removed from the incubator and immediately placed on ice. The medium was removed and immediately processed for EV purification. Cells were washed with ice-cold PBS and lysed in ice-cold 1X RIPA buffer (Cell Signaling 9806S) with 1X Protease inhibitors (Santa Cruz sc-29131). Cells were incubated on ice for 10min then transferred to tubes using cell scrapers. Lysates were cleared at 20,000xg for 10min at 4°C. The pellet was discarded. Total protein was measured using Pierce BCA Protein Assay Kit according to the manufacturer’s microplate protocol (Thermo Fisher 23225). A final concentration of 1X Laemmli sample buffer (Bio-Rad 161-0747) with 10% beta-mercaptoethanol (BME) (Bio-Rad 161-0710) was added to 15µg of total protein.

EV and mixed samples were vortexed for 30s and incubated for 10min at room temperature in 1X RIPA buffer (Cell Signaling 9806S) and 1X Protease inhibitors (Santa Cruz sc-29131) to lyse EVs. Total protein was measured using Pierce BCA Protein Assay Kit according to the manufacture’s microplate protocol (Thermo Fisher 23225). 1X Laemmli sample buffer (Bio-Rad 161-0747) was added to 1µg of total protein.

### Immunoprecipitation

Mixed and protein fractions were pre-cleared of IgG from FBS with 100µL of protein G coated magnetic Dynabeads (Thermo Fisher 10003D Lot# 00715594) overnight at 4°C with rotation. 50µL of protein G beads were coated 2ug of anti-AGO2 antibody (Sigma SAB4200085 Lot# 0000089486) according to the manufacturer’s instructions. The clearing protein G beads were removed from the sample and replaced with the anti-AGO2 coated beads overnight at 4°C with rotation. The beads were washed three times with 1X PBST and resuspended in 1X Laemmli sample buffer 10% BME (Bio-Rad 161-0747).

### Western Blot Analysis

Samples were heated to 95°C for 5min then separated alongside a spectra multi-color Ladder (Thermo Fisher 26634) through a 4-15% Tris-Glycine extended Stain-Free gel (Bio-Rad 5678085), at 100V for 1.5hrs using 1X Tris-Glycine SDS buffer (Bio-Rad 161-074). Gels were imaged with an EZ DocGel Stain-free imaging system (Bio-Rad 170-8274). Proteins were then transferred to a methanol activated (10s) PVDF membrane (Bio-Rad 1620177) at 100V for 1hr in 1X Tris-Glycine buffer (Bio-Rad 161-0734) at 4°C. The PVDF membrane was blocked for 1hr. in blocking buffer (1XPBS (Gibco 14190-144), 0.05% Tween20 (Sigma-Aldrich 274348500), 5% blotting-Grade Blocker (Bio-Rad170-6404)). The membrane was incubated with primary antibodies anti-CD63 (BD 556019 Lot#7341913) diluted 1:1000 and anti-CD81(sc-7637 Lot#C2318) diluted 1:500, anti-CD9 (BioLegend 312102 Lot#B351275), ant-TSG101 (abcam ab228013 Lot#GR3306738-8) diluted 1:500, anti-Syntenin (abcam ab133267 Lot# GR3375272-1) diluted 1:500, anti-GM130 (ab52649 Lot#GR3427322-2, anti-Calnexin (ab22595) diluted 1:1,000, anti-Albumin (abcam ab28405 Lot#GR3367930-2) diluted 1:1000, or anti Hif-1a (Cayman 10006421) diluted 1:200, anti-AGO2 (Sigma SAB4200085 Lot# 0000089486) diluted 1:1,000 and anti-beta-actin (Sigma A1978) diluted 1:10,000 in blocking buffer overnight at 4°C with rotation. The membrane was washed 3X for 5min with rotation in blocking buffer. Secondary anti-mouse-HRP (Santacruz sc-516102 Lot#c1419) was diluted 1: 10,000 or anti-rabbit-HRP (Dako P0448) 1:1000 in blocking buffer and incubated for 1hr at room temperature. Membranes were washed 3X in blocking buffer then 2X in 1XPBS 0.05% Tween20. Membranes were then incubated with Super Signal Chemiluminescent Substrate (Thermo 34580) for 5min with gentle rotation and imaged by iBright FL1000 (Invitrogen). For HIF-1a, band intensity in normoxia was normalized to hypoxia using Image J software.

### RNA Isolation

Total RNA was extracted from cells using mirVana miRNA isolation kit (Ambion cat#AM1560 following the manufacturer’s protocol for adherent cells. Total RNA was extracted from size-exclusion chromatography (SEC) EV and protein fractions using miRNeasy serum/plasma kit (Qiagen 1071073 Lot#160020206) after adding 1.0×10^6 copies/µL of cel-miR-39 exogenous spike-in control (Qiagen 219610 Lot#157036035), according to the manufacturer’s instructions. RNA concentration and purity were measured by NanoDrop (Thermo Fisher).

### Small RNA Library Construction and Analysis

The ligation-independent Capture and Amplification by Tailing and Switching (CATS) small RNA-seq method was used to profile cellular small RNA. Libraries were constructed with CATS RNA-seq Kit (Diagenode C05010041 Lot#4) following the manufacturer’s instructions. Library primer dimers were removed by AMPureXP beads (Beckman). Quality control was assessed by bioanalyzer high sensitivity assay (Agilent). Libraries were then size selected between 160-180bp by BluePippin (Sage Science). Prior to sequencing, libraries were spiked with 20% PhiX Control v3 (Illumina 1501766) then run using the NovaSeq Illumina sequencing platform by the Johns Hopkins Microarray and Deep Sequencing Core. Sequencing quality control was performed using FASTQC (Babraham Bioinformatics) followed by adapter and PolyA trimming with Cutadapt 1.17. The reads were sequentially aligned to reference transcriptomes rRNA, tRNA, RN7S, snRNA, snoRNA, scaRNA, VT-RNA and Y-RNA. All reads that did not map to the aforementioned RNAs were aligned to custom curated pre-miRNA, protein coding mRNA and long non-coding RNAs (lncRNAs) references. All low expressed miRNA, lncRNA and mRNA reads were filtered out (average number of reads for a gene in all samples >=5). For each sample, all reads that did not map to the human transcriptome were aligned to the hg38 genome reference (rest_hg38) and thus corresponded to reads aligned to introns and intergenic regions combined. Total hg38 reads were used to calculate log2[counts-per-million+1] for each gene. The log-fold-change (LFC), a log2[cpm+1] difference between conditions was used as a final metric to determine relative abundance. See supplementary section for exact bash scripts. The differential expression analysis was done using edgeR/limma pipeline as described in ^54^.

### RT-qPCR

Two step RT-qPCR was performed for miRNA analysis starting with 2µL of RNA input from cells, EVs, and Ex-protein fractions using TaqMan microRNA Reverse transcription kit (ABI 4366597 Lot#00636931), TaqMan miRNA stem-loop RT primers/qPCR primer probe set (ABI 4427975): (cel-miR-39 ID# 000200 Lot#P180110-003B10, miR-16 ID# 000391 Lot#P171018-000H05, miR-210-3p ID#000512), and TaqMan Universal Master Mix II, no UNG (ABI 4440040 Lot#1802074) as described in manufacturer’s protocol. For mRNA analysis, High capacity cDNA synthesis kit (ABI cat#4368813), TaqMan qPCR primer probe set ABI: (HIF-1α ID#Hs00153153_m1, GAPDH ID# Hs02786624_g1, Beta-Actin ID# Hs01060665_g1) and TaqMan Universal Master Mix II, no UNG was used following kit protocol. qPCR was run on a CFX96 Real-time System (Bio-Rad). miR-210 was normalized to miR-16 and cel-miR-39 using the 2^−ΔΔCT^ method. HiF-1α was normalized to the average of GAPDH and Beta-actin Pooled by 2^−ΔΔCT^.

### Data Availability

All relevant experimental details and data were submitted to the EV-TRACK knowledgebase (EV-TRACK ID: EV200090)^55^ and Gene Expression Omnibus (GEO), accession numbers GSE207677 and GSE212703.

## RESULTS

### miR-210-3p expression induced by HIF-1α stabilization in SW1736

To confirm the inducible expression of miR-210-3p in response to HIF-1α stabilization in hypoxia, SW1736 (ATC) cells were cultured at 21% O_2_ (normoxia) or 2% O_2_ (hypoxia) for 24, 48, 72, 96, and 120hrs. Increased HIF-1α protein levels in hypoxia were observed at each time point; the greatest increase compared with normoxic controls were at 24 and 48hrs (Figure1.B). Concurrent with increased HIF-1α protein levels, miR-210-3p expression increased at each hypoxic time point, with the highest expression levels also at 24hrs (~9-fold) and 48hrs (~16-fold) relative to normoxia (Figure1.C). Both HIF-1α and miR-210-3p levels slightly declined by 72hrs and thereafter (Figure1.B-C). This is consistent with our observation and previous reports of decreased levels of HIF-1α protein due to mRNA instability in prolonged hypoxia (Figure1.D)^56–58^ as well as negative regulation of HIF-1α by miR-210 in a study of human T-cells^59^.

**Figure 1.**
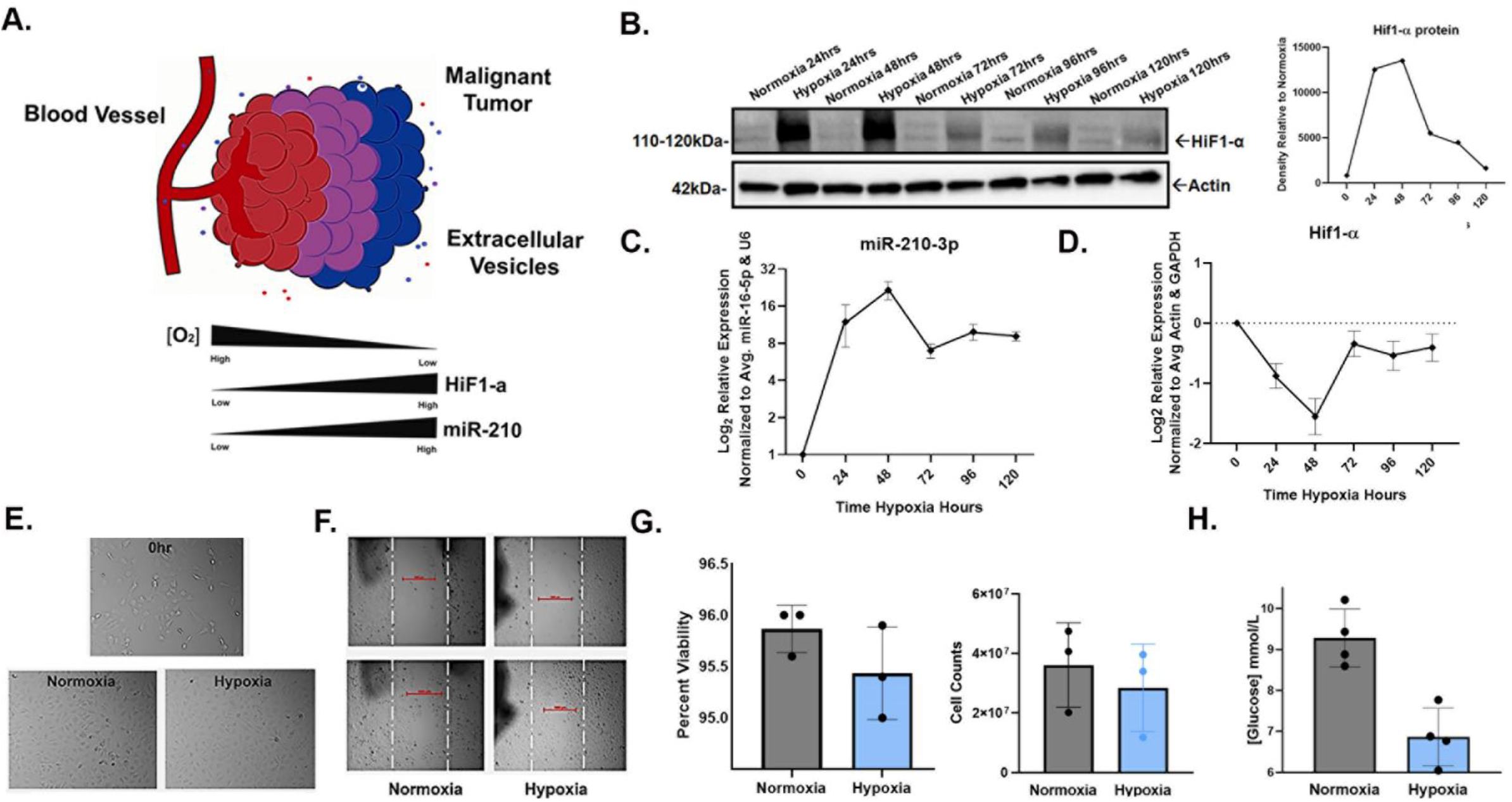
HIF1-α-induced miR-210-3p expression during hypoxic culture of SW1736 cells. **A**. Cartoon illustration of hypoxic malignant tumor. In low oxygen conditions, miR-210 expression is regulated by HIF. **B**. HIF1-α stabilization verified by Western blot (WB) and densitometry analysis after 24,48,72,96, and 120hrs of hypoxia (2% O_2_) or normoxia (21% O_2_). Actin was used as a loading control. **C**. RT-qPCR analysis of mir-210-3p expression in hypoxia vs. normoxia at 24, 48, 72, 96, and 120hrs of hypoxia relative to normoxia. Error bars represent the standard deviation of 3 biological repeats. **D**. RT-qPCR analysis of HIF-α expression relative to normoxia **E**. Images of SW1736 cells before and after 72 hours of normoxia or hypoxia. **F**. Wound Healing assay of SW1736. **G**. Percent viability of SW1736 cells after 72hrs of normoxia or hypoxia. SW1736 cell counts after 72hrs of normoxia or hypoxia. **H**. SW1736 cell culture media glucose concentration after 72hrs of normoxia or hypoxia.

Additionally, cell morphology, viability and glucose consumption were compared in hypoxia vs. normoxia. Cells from each condition were visibly comparable (Figure1 E-F) and shared a similar viability, but cell counts were lower in hypoxia (Figure1.G). Additionally, glucose consumption significantly increased in hypoxia (Figure1.H). Taken together, our results confirm the positive correlation between miR-210-3p expression and HIF1-α protein levels in SW1736.

### Hypoxia induces miR-210 more strongly in ATC compared with PTC lines

Small RNA sequencing was performed on ATC cell lines SW1736 and C643 and PTC cell lines BCPAP and TPC-1, all grown under normoxic culture conditions. We identified 28 miRNAs that were differentially expressed (fold-change >2, p-value <0.05) between normoxic ATC and PTC lines (Figure2.A). Under normoxia, miR-210 was ~5-fold less abundant in ATC lines versus PTC lines (Figure2.A). qPCR validation confirmed this result for normoxia (Figure2.B). However, in hypoxia, ATC miR-210 expression reached levels comparable to or greater than those in PTC cells (Figure2.B). A time course assay examining points from 0-48hrs of hypoxia revealed that ATC cell lines showed a higher fold change, >2-fold, of miR-210-3p expression compared with the PTC cell lines (Figure2.C). Therefore, although basal (normoxic) levels of miR-210 are lower in ATC than PTC lines, the miRNA is more strongly up-regulated by hypoxia in ATC lines.

**Figure 2.**
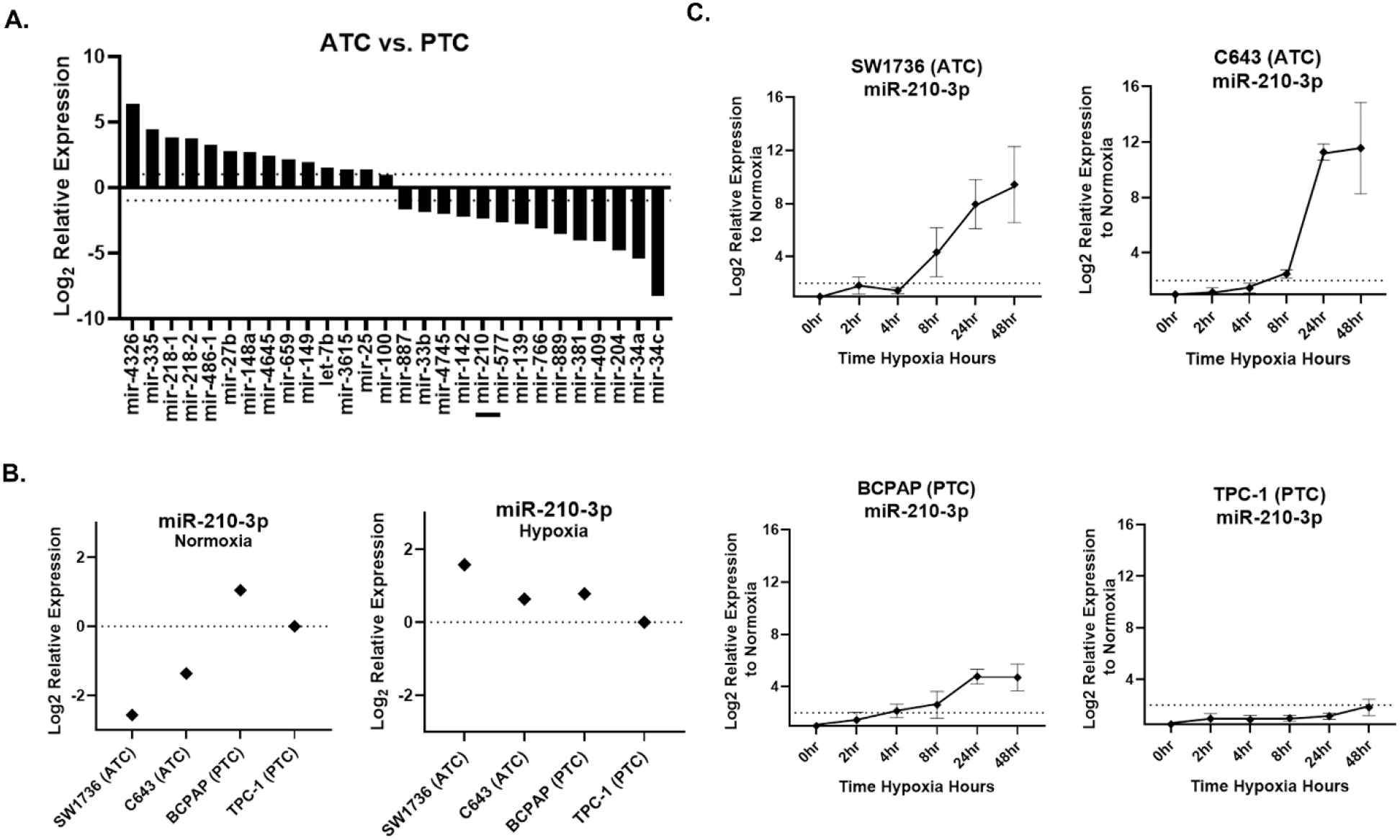
miR-210 expression in ATC and PTC cell lines. **A**. RNA-seq analysis of differentially expressed pre-miRNAs between anaplastic thyroid cancer (ATC) cell lines SW1736 & C643 and papillary thyroid cancer (PTC) cell lines BCPAP and TPC-1. Log_2_ fold change>+/-1. q value < 0.05 n=3. Bar included to highlight miR-210. **B**. RT-qPCR of miR-210 basal expression and hypoxia (2%O_2_) induced expression in ATC vs. PTC cell lines. **C**. ATC vs. PTC qPCR analysis of miR-210-3p expression during hypoxia time course study at 0,2,4,8,24,48hrs hypoxia.

### miR-210-3p is the primary hypoxia-responsive miRNA in SW1736 cells

To examine how hypoxia might impact the expression of additional miRNAs in SW1736, small RNA-sequencing was performed to assess miRNA differences in normoxia vs hypoxia at 72 hours. Although miR-210-3p was up-regulated in hypoxia (~6-fold, p<0.05), no other miRNAs were significantly differentially expressed greater than 2-fold, p<0.05 (Figure3. A). We also inspected the differential expression of protein-coding mRNA in response to hypoxia in SW1736 by RNA-seq analysis. 8 transcripts were significantly up-regulated >2-fold, p<0.05 (Figure3.B-C). Surprisingly, no genes were significantly down-regulated. Hierarchical clustering emphasized coordinated responses to hypoxia.

**Figure 3.**
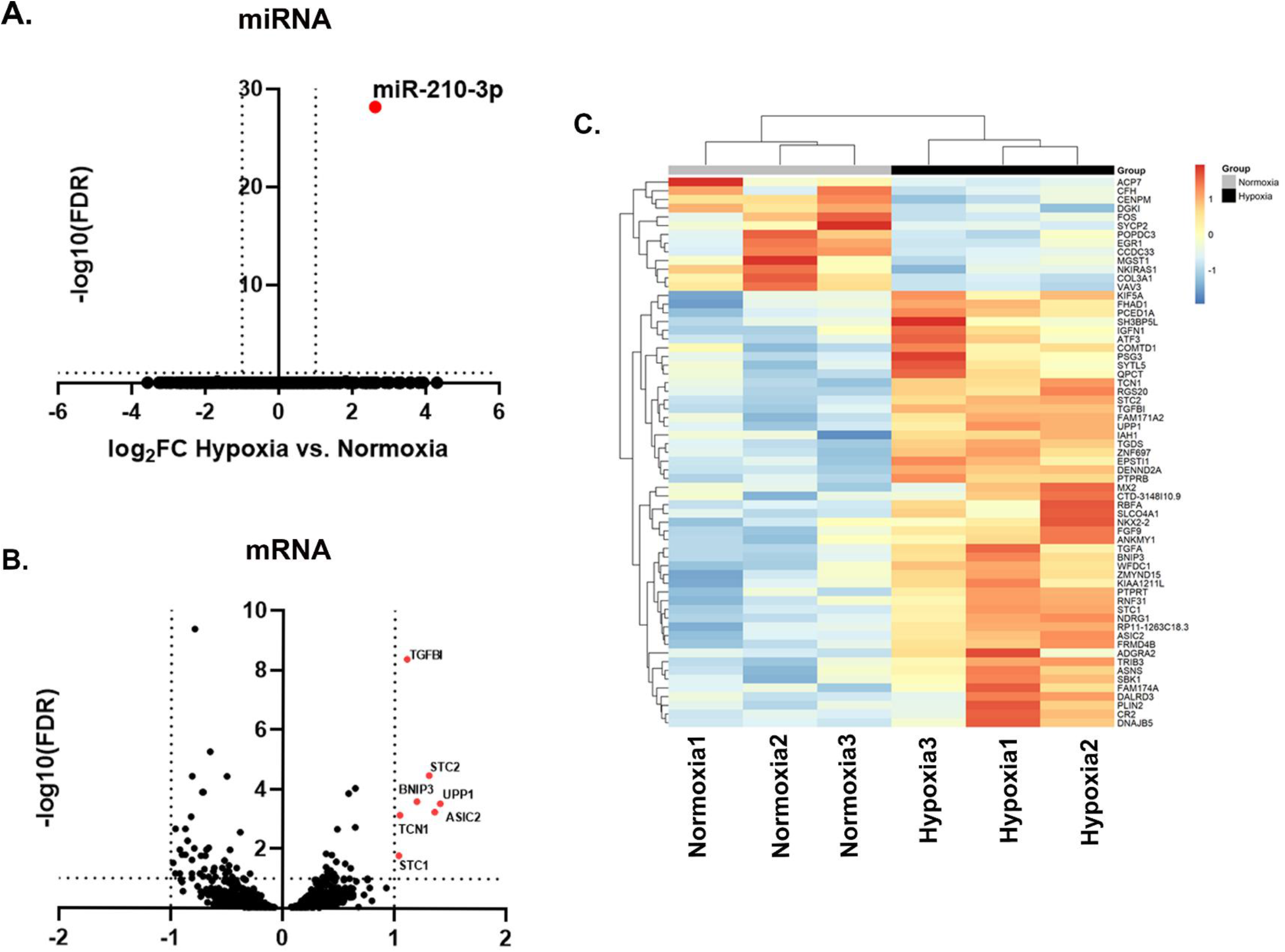
SW1736 cellular miRNA expression in hypoxia at 72hrs. **A**. Differential expression of miR-210-3p in hypoxia vs. normoxia shown as a volcano plot of false discovery rate (FDR) and log_2_ fold change (FC). **B**. Log_2_ FC of differentially expressed mRNAs with base Mean > 10, log2Fold Change > +/-1 and raw p value < 0.05 and adjusted p value < 0.05. **C**. Heatmap of the normalized read counts for all differentially expressed mRNAs with base Mean > 10, log2Fold Change > +/-1 and raw p value < 0.05 (n=63 in total).

### Increase of miR-210-3p release from SW1736 cells in hypoxia

Cells and conditioned cell culture media (CCM) were collected after 72hrs of normoxic and hypoxic culture, and CCM was fractionated by size-exclusion chromatography (SEC) to obtain fractions enriched in extracellular vesicles (EVs), intermediate mixed EVs and proteins, and extracellular proteins. Fractions were characterized per MISEV2018 guidelines^60^. Presence of EVs was verified by Western blot (WB) using EV-rich markers CD63, CD9, CD81, TSG101, and Syntenin, which were detected in EV-rich fractions but not in the later protein fractions (Figure4.B). The depletion of cellular markers such as GM130 and calnexin in extracellular fractions was confirmed relative to the cell source. (Figure4.B). Single-particle interferometric reflectance imaging sensing (SP-IRIS) further confirmed EV-rich markers CD63, CD9, and CD81 relative to MIgG control (Figure4.C). Fractions were imaged by transmission electron microscopy (TEM), revealing that EVs ranged in diameter from ~50-300nm in both normoxia and hypoxia in pooled EV-rich SEC fractions; EVs were not observed in protein-rich fractions (Figure5.C). We also assessed particle concentration and size by nano-flow cytometry (NFCM) (Figure4.E), finding no significant differences. qPCR for miR-210-3p showed that miR-210-3p was released from cells in all fractions during normoxia, and that this release increased, also across all extracellular fractions, by >2-fold in hypoxia (Figure4.F). Finally, immunoprecipitation confirmed the presence of miRNA carrier AGO2 in both mixed and protein fractions (Figure4.G)

**Figure 4.**
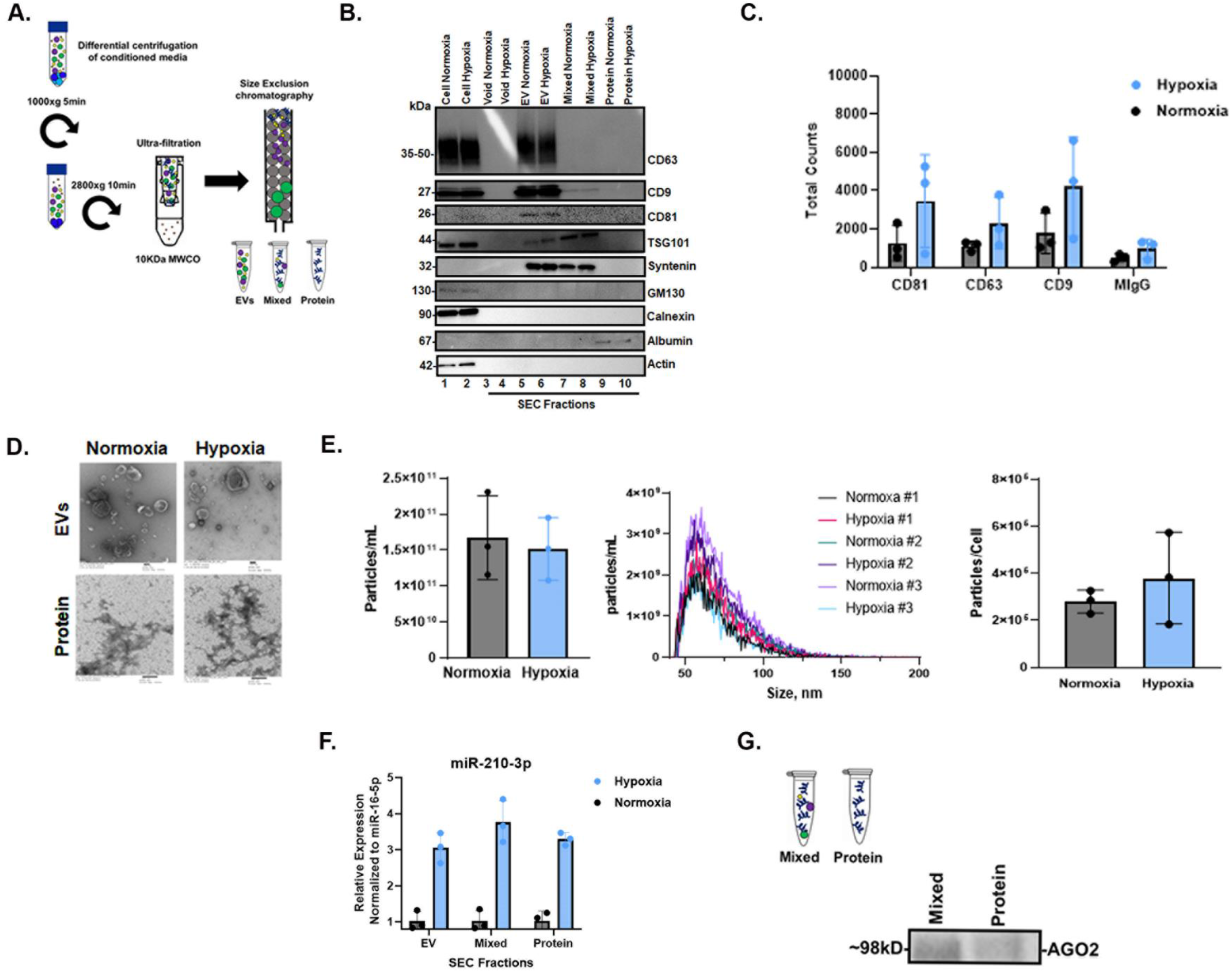
Characterization of cellular and extracellular fractions and AGO2 detection in hypoxic SW1736 culture. **A**. Workflow diagram schematic of EV and protein separation from SW1736 cell culture media (CCM) by differential centrifugation and size-exclusion chromatography (SEC). Three independent experiments were performed. **B**. Western blot analysis of tetraspanins CD63 and CD81 expression in SW1736 cell lysates and enrichment in hypoxic and normoxic SEC fractions after 72hrs. **C**. SP-IRIS detection of tetraspanins CD81, CD63, and CD9 in hypoxic and normoxic EV enriched SEC fractions after 72hrs.**D**. Transmission electron micrograph (EM) of hypoxic and normoxic EV and protein SEC fractions after 72hrs. **E**. SEC EV fraction particle counts per cell from normoxia and hypoxia. Data points represent 3 independent experiments by nano-flow cytometry. **F**. miR-210-3p SEC pooled EV and protein qPCR analysis after 72hrs hypoxia or normoxia. Derived from 100mLs of culture media. Data points represent 3 independent experiments. **G**. AGO2 detection in SEC mixed and protein fractions by immunoprecipitation followed by Western blot.

## DISCUSSION

Highly aggressive tumors, which are often hypoxic, are frequently resistant to oxygen-dependent treatments such as chemotherapy and radiation. Thus, several direct and indirect methods to assess tumor hypoxia for targeted therapy, including miRNA expression, have been explored^61–65^. However, to our knowledge, miRNAs have not been extensively studied in hypoxic vs. non-hypoxic regions within ATC tumors. In this work we examined miR-210, a direct HIF-1 target, as a potential marker of hypoxia in ATC^66,67^. Our results demonstrate that precursor miR-210 is down-regulated in ATC cell lines compared with PTC cells, but it is more robustly up-regulated in ATC in response to hypoxia. Additionally, we show that miR-210-3p can be detected extracellularly and is associated with EVs and extracellular AGO2.

miR-210 has paradoxically been described as both oncogenic and as a tumor suppressor in many studies ^42,68,69,70^. It is believed to be involved in cell cycle regulation, metabolism, and angiogenesis^71,72^. However, the exact role of miR-210-3p in tumorigenesis remains elusive. In this study, we also examined the differential expression of mRNA, but no down-regulated transcripts were identified. Polysome profiling by proteomic analysis of ATC cell lines in hypoxia may be required to identify putative miR-210-3p targets in this cell type, as the use of a miR-210-3p mimic may result in off-target gene down-regulation.

It was interesting to observe lower basal levels of miR-210-3p in ATC cell lines SW1736 and C643, which are de-differentiated, compared with poorly and well-differentiated PTC cell lines BCPAP and TPC-1, respectively. This finding suggests that miR-210 might also be linked to tumor differentiation. It is believed that ATC can originate from PTC^73–75^; therefore, it is possible that miR-210 expression is impacted during de-differentiation. However, because miR-210 down-regulation was observed at both the precursor and mature level, it is unclear if the basal level expression differences are directly related to HIF-1 or due to miR-210 genetic or epigenetic alterations. In either case, lower basal expression levels make miR-210 induction more pronounced in hypoxia; hence, miR-210-3p may be a better marker of hypoxia in ATC compared with PTC.

We hypothesized that miR-210-3p might also be detected extracellularly in hypoxia, as miRNAs are remarkably stable in biological fluids such as blood plasma and are promising non-invasive biomarkers^29,32,34,35^. This is ascribed to their association with RNA carriers such as EVs and RNA-binding proteins, which offer protection from degradation^34,76^. In our study, we observed increased extracellular miR-210-3p in conditioned cell culture medium in hypoxia after at least 48hrs, which further increased after 72hrs. This suggests that, although cellular levels of miR-210 increase almost immediately in response to hypoxia, additional time is needed for the newly produced miRNAs to pass into the extracellular space and be detected there.

Apart from miR-210-3p, we documented upregulation of various small RNA fragments mapped to several protein coding transcripts including TGFBI, STC2, BNIP3, UPP1, ASIC2, TCN1, and STC1. The mechanism of formation and the biological role of these mRNA-aligned small RNAs remains unknown, however, we suggest that they could be used as promising biomarkers for hypoxia in addition to miR-210-3p.

**In summary**, our observations, albeit limited to *in vitro* studies, suggest that miR-210-3p might be a suitable miRNA marker of hypoxia in ATC.

